# Enhanced Maturity and Functionality of Vascularized Human Liver Organoids through 3D Bioprinting and Pillar Plate Culture

**DOI:** 10.1101/2024.08.21.608997

**Authors:** Vinod Kumar Reddy Lekkala, Sunil Shrestha, Ayah Al Qaryoute, Sanchi Dhinoja, Prabha Acharya, Abida Raheem, Pudur Jagadeeswaran, Moo-Yeal Lee

**Affiliations:** Department of Biomedical Engineering, University of North Texas, Denton, Texas, USA; Department of Biological Sciences, University of North Texas, Denton, TX, USA; Bioprinting Laboratories Inc., Dallas, Texas, USA

**Keywords:** Vascular human liver organoids, pillar plate, deep well plate, coagulation factor secretion, enhanced organoid maturity

## Abstract

Liver tissues, composed of hepatocytes, cholangiocytes, stellate cells, Kupffer cells, and sinusoidal endothelial cells, are differentiated from endodermal and mesodermal germ layers. By mimicking the developmental process of the liver, various differentiation protocols have been published to generate human liver organoids (HLOs) *in vitro* using induced pluripotent stem cells (iPSCs). However, HLOs derived solely from the endodermal germ layer often encounter technical hurdles, such as insufficient maturity and functionality, limiting their utility for disease modeling and hepatotoxicity assays. To overcome this, we separately differentiated EpCAM^+^ endodermal progenitor cells (EPCs) and mesoderm-derived vascular progenitor cells (VPCs) from the same human iPSC line. These cells were then mixed in BME-2 matrix and concurrently differentiated into vascular human liver organoids (vHLOs). Remarkably, vHLOs exhibited significantly higher maturity than vasculature-free HLOs, as demonstrated by increased coagulation factor secretion, albumin secretion, drug-metabolizing enzyme (DME) expression, and bile acid transportation. To enhance assay throughput and miniaturize vHLO culture, we 3D bioprinted expandable HLOs (eHLOs) in BME-2 matrix on a pillar plate platform derived from EPCs and VPCs and compared with HLOs derived from endoderm alone. Compared to HLOs cultured in a 50 μL BME-2 matrix dome in a 24-well plate, vHLOs cultured on the pillar plate exhibited superior maturity, likely due to enhanced nutrient and signaling molecule diffusion. The integration of physiologically relevant patterned liver organoids with the unique pillar plate platform enhanced the capabilities for high-throughput screening and disease modeling.

## Introduction

The liver’s complex physiological functions are sustained by intricate intracellular communication among cells derived from the endodermal, mesodermal, and ectodermal germ layers. Although recent advancements in liver organoid technology are promising, they encounter limitations due to the diverse cellular composition necessitated by these distinct germ layers. Liver organoids, which are primarily composed of hepatocytes and cholangiocytes from the endodermal layer, frequently face challenges such as insufficient cellular complexity, maturity, and incomplete functionality. These issues limit their utility in disease modeling and drug testing applications.

Vascular progenitor cells originating from the mesodermal germ layer that differentiate into key components of the vascular system, including endothelial cells and pericytes. These cells are essential for forming blood vessels, ensuring proper nutrient and oxygen delivery within tissue. By integrating mesoderm-derived vascular cells with endoderm-derived liver organoids, the physiological relevance of the liver organoids could be significantly enhanced by improved vascularization. This integration is crucial for supporting organoid growth and functionality, addressing the current limitations in liver organoid development. There have been several vascular liver models, such as multicellular liver models, developed by combining heterogeneous cell sources of endothelial cells and liver-associated cell types^1–6^. Co-cultures of primary liver cell types also struggle to achieve the same level of *in vivo* cell and tissue function. In addition, reproducibility, scale-up production, and user-friendliness of primary liver cell types for preclinical evaluations remain technically challenging.

Recently, significant attention has been directed towards human liver organoids (HLOs) derived from induced pluripotent stem cells (iPSCs) due to their complexity, multicellularity, and functional resemblance to the native liver, including their ability to secrete drug-metabolizing enzymes (DMEs)^7,8^. This advancement opens new avenues for studying liver biology, disease modeling, and pharmaceutical applications. The combination of germ layers such as mesoderm and endoderm have been shown to enhance the complexity and maturity of tissue models, including tumor organoids, gastrointestinal organoids, and pulmonary organoids^9–11^. In this study we generated vascular human liver organoids (vHLOs) by combining EpCAM^+^ endodermal progenitor cells (EPCs) and mesoderm-derived vascular progenitor cells (VPCs) from the same human iPSC line in BME-2 matrix. In addition, vHLOs were generated on a high-throughput pillar plate platform by 3D bioprinting of expandable HLOs (eHLOs) derived from EPCs and VPCs.

Liver organoids on a high-throughput screening (HTS) platform could offer unparalleled opportunities to recapitulate human tissue architecture and function *in vitro* for studying complex biological processes, disease mechanisms, and drug responses. Several engineering approaches have recently developed as alternatives to animal testing, which could potentially simulate organ-level functions *in vitro*^12^. Culturing human organoids on novel culture platforms could recapitulate critical organ features, potentially reducing or replacing animal testing while reducing costs and time. In this study we made advancements in improving the maturity and functionality of liver organoids by co-differentiation of EPCs and VPCs and 3D bioprinting of eHLOs on a pillar plate. The incorporation of mesodermal vascular components into endodermal liver organoid culture enhanced liver organoid maturity and function, thereby increasing their physiological relevance.

## Materials and Methods

### Fabrication of pillar and deep well plates

A 36-pillar plate with sidewalls and slits (36PillarPlate) and a clear-bottom 384-deep well plate (384DeepWellPlate) were manufactured by the injection molding of polystyrene (**Figure 1**) for cell printing and organoid culture (Bioprinting Laboratories Inc., Dallas, TX, USA)^13,14^. The 36PillarPlate contains a 6 x 6 array of pillars (4.5 mm pillar-to-pillar distance, 11.6 mm pillar height, and 2.5 mm outer and 1.5 mm inner diameter of pillars). The 384DeepWellPlate has a 16 x 24 array of deep wells (3.5, 3.5, and 14.7 mm well width, length and depth, and 4.5 mm well-to-well distance) for cell culture. For organoid culture, the pillar plate with cells encapsulated in BME-2 matrix was inverted and sandwiched onto the 384DeepWellPlate.

**Figure 1.**
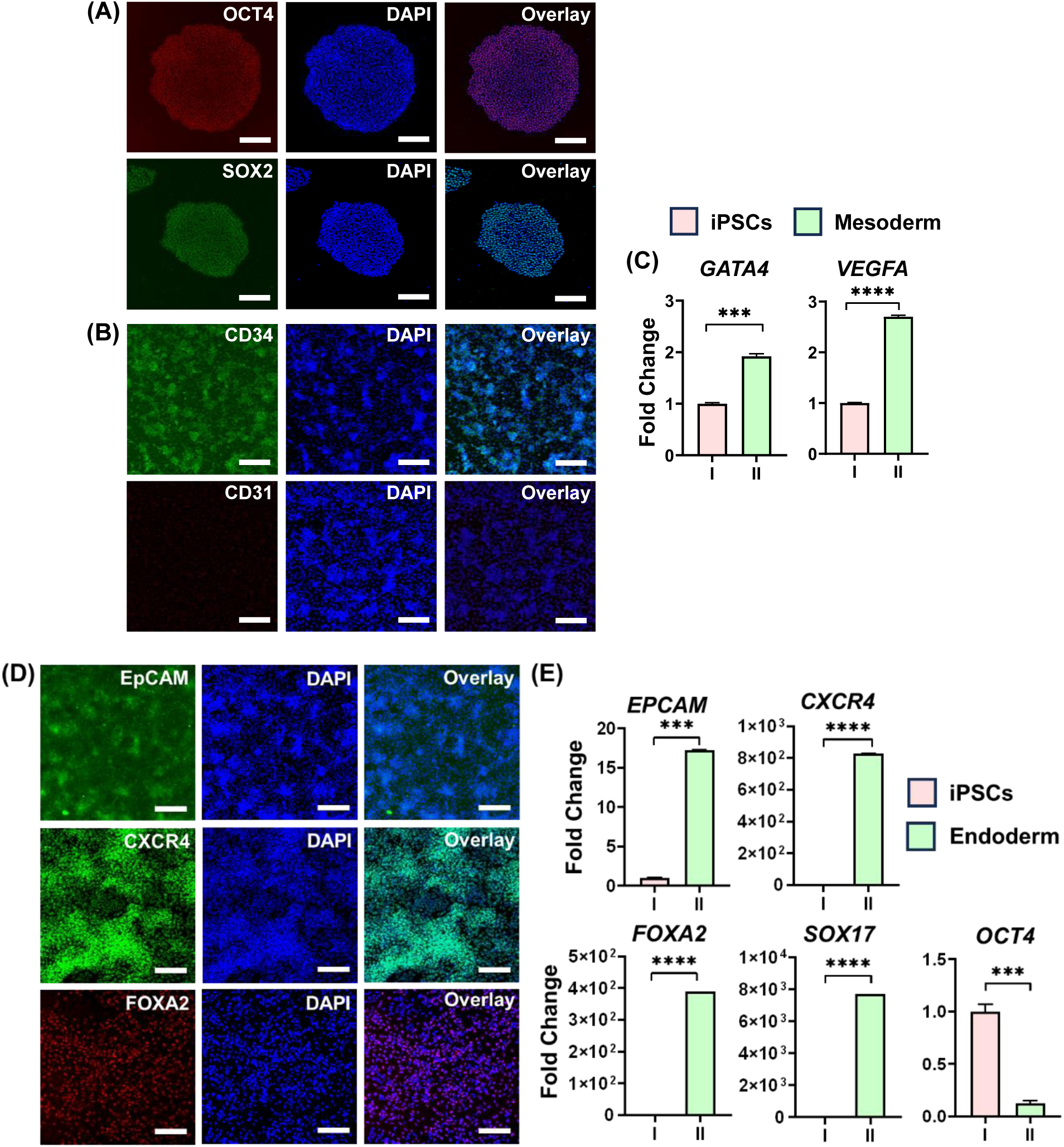
Generation of endodermal progenitor cells (EPCs) and mesodermal vascular progenitor cells (VPCs) from human iPSCs. **(A)** Immunofluorescence staining of human iPSCs for OCT4 and SOX2 stem cell markers. Scale bars: 200 µm. **(B)** Immunofluorescence staining of vascular progenitor cells after 3 days of iPSC differentiation for CD34 endothelial progenitor marker and CD31 endothelial marker. Scale bars: 200 µm. **(C)** The expression level of mesoderm genes including *GATA4* mesodermal cells as well as *VEGFA* endothelial cells. n = 3. The significance of gene expression difference was analyzed by Student’s t test. **(D)** Immunofluorescence staining of endodermal progenitor cells after 5 days of iPSC differentiation for EpCAM endodermal progenitor cells as well s CXCR4 and FOXA2 endodermal markers. Scale bars: 200 µm. **(E)** The expression level of endoderm genes including *EPCAM* endodermal progenitor cells, *CXCR4*, *FOXA2*, and *SOX17* endodermal cells, and *OCT4* stem cells. n = 3. The significance of gene expression difference was analyzed by Student’s t test.

### Differentiation of iPSCs into EpCAM^+^ endodermal progenitor cells (EPCs)

A male human iPSC line, EDi029-A (Cedar Sinai Biomanufacturing Center, USA), was maintained in an iMatrix-511 silk (Elixirgen Scientific; NI511) coated 6-well plate using mTeSR^TM^ Plus medium (Stemcell Technologies; 100-0276). The differentiation of iPSCs into EPCs was performed using a previously published protocol^15^, with some modifications. Briefly, at 70 - 80% confluency, iPSCs were harvested using Accutase (Gibco; A1110501), seeded in a 6-well plate at a cell density of 0.3 x 10^6^ cells/well and cultured using mTESR^TM^ Plus medium supplemented with 5 µM Y27632 Rho-kinase (ROCK) inhibitor (Tocris; 1254) for the first 24 hours. After 24 hours of culture, the iPSCs were differentiated into endodermal progenitor cells using RPMI 1640 (Gibco; 22400089) supplemented with 100 ng/mL activin A (Tocris; 338-AC), 50 ng/mL recombinant human Wnt3a (R&D Systems; 5036-WN), 5 ng/mL R-Spondin1 (R-spo1; R&D systems; 4645-RS/CF), and 1% B27 supplement (Gibco; 17504) for 5 days. The cell culture medium was changed every day.

### Differentiation of mesodermal vascular progenitor cells (VPCs)

The differentiation of EDi029-A iPSC line into vascular mesodermal cells was performed using a previously published protocol^16^, with some modifications. Briefly, the iPSC line was maintained in the iMatrix-511 silk coated 6-well plate using mTeSR^TM^ Plus medium. At 70 - 80% confluency, the mTeSR Plus medium was replaced with a mesoderm layer induction medium containing 5 µM CHIR99021 (R&D Systems; 4423), 25 ng/mL bone morphogenetic protein 4 (BMP4; Tocris; 314-BP), 1% GlutaMAX^TM^ (Gibco; 35050), and 1% B27 in DMEM/F12 (Gibco; 12634) for 3 days.

### Generation of expandable human liver organoids (eHLOs) in a 24-well plate

After differentiation, EPCs and VPCs were detached using Accutase, pipetted several times to make single cells, and centrifuged at 200 rcf for 3 minutes. The EPCs and VPCs were mixed in the ratio of 4:1, suspended in Cultrex basement membrane extract, type 2 (BME-2; R&D Systems; 3532-010-02), and dispensed in a 24-well plate to form 50 µL BME-2 domes for culture. After gelation of BME-2 at 37°C for 10 - 12 minutes, the cell population was differentiated in advanced DMEM/F12 (Gibco; 12634) supplemented with 1% B27 supplement (Gibco), 1.25 mM N-acetyl-L-cysteine (Sigma; A9165), 10 nM recombinant human (Leu^15^)-gastrin I (Sigma; G9145), 100 ng/mL recombinant human fibroblast growth factor 10 (FGF10; Peprotech; 100-26), 25 ng/mL recombinant human hepatocyte growth factor (HGF; Peprotech; 100-39), 10 mM nicotinamide (Sigma; N0636), 5 μM A83-01 (R&D Systems; 2939), 10 μM forskolin (R&D Systems; 1099), 100 ng/mL recombinant human VEGF (R&D; 293-VE-010/CF), and 10% R-spo1 conditioned medium prepared by culturing Cultrex HA-R-Spondin1-Fc 293T cells (R&D Systems; 3710-001-01) to generate premature, expandable liver organoids. For the first 3 days of culture, the medium was supplemented with 25 ng/mL Noggin (R&D Systems; 6057-NG-025/CF), 10 μM Y27632, and 30% Wnt-conditioned medium prepared by culturing L Wnt-3A cells (ATCC; CRL-2647). From day 4, the medium was changed every 3 days. After 10 - 14 days of culture, the expandable organoids grew to 300 - 500 µm in diameter, which were collected from BME-2 dome through mechanical dissociation using 1 mL pipette. Passaging of the expandable organoids was performed at a 1:4 split ratio in BME-2 every 6 - 8 days when the desired confluency was reached.

### Maturation of vascular human liver organoids (vHLOs) in a 24-well plate

The dissociated cell clumps from eHLOs were mixed with undiluted BME-2 and dispensed at 50 μL BME-2 dome/well in a 24-well plate for maturation. The cell clumps in BME-2 dome were cultured in advanced DMEM/F12 supplemented with 1% B27, 10 nM gastrin, 25 ng/mL HGF, 25 ng/mL human active BMP7 recombinant protein (Gibco; PHC7204), 100 ng/mL recombinant human FGF19 (Peprotech; 100-32), 500 nM A83-01, 10 μM notch inhibitor DAPT (Sigma; D5942), and 30 μM dexamethasone (Dex; Sigma; D4902). This medium was replaced every 3 days for 20 days for maturation. For functional assays, the medium was collected on day 20 after 3 days of culture and stored in a -80° freezer until use for analysis. All the experiments were carried out with eHLOs at passage numbers 3 - 6 (**Supplementary Figure 3**).

### Microarray 3D bioprinting of eHLOs in BME-2 on the pillar plate for the differentiation of vHLOs

The cell clumps obtained from physical dissociation of eHLOs were suspended in undiluted BME-2, and 5 μL spot of the cell-hydrogel suspension was printed on the 36PillarPlate using ASFA^TM^ Spotter V6 (MBD Co., Ltd., South Korea). After BME-2 gelation at 37°C for 10 - 12 minutes, the 36PillarPlate with cell clumps was sandwiched onto a complementary 384DeepWellPlate containing 80 µL of the maturation medium in each well. This medium was replaced every 3 days for 20 days.

### Gene expression analysis *via* RT-qPCR

HLOs in BME-2 were either collected manually by pipetting in cold PBS^-/-^ or isolated from BME-2 using Cultrex organoid harvesting solution (R&D Systems; 3700-100-01) according to the manufacturer’s protocol. Total RNA was isolated from harvested cells by using the RNeasy Plus Mini Kit (Qiagen; 74134) following the manufacturer’s recommended protocol. cDNA was synthesized from 1 µg of RNA using the high-capacity cDNA reverse transcription kit protocol (Applied Biosystems; 4368814). Real time PCR was performed using PowerTrack^TM^ SYBR green master mix (Applied Biosystems; A46110) and forward/reverse primers from IDT Technology in the QuantStudio™ 5 Real-Time PCR System (Applied Biosystems; A28574). The primers used are listed in **Supplementary Table 1**. The expression level of target genes was normalized against the housekeeping gene, glyceraldehyde 3-phosphate dehydrogenase (GAPDH).

### Whole mount immunofluorescence staining

The immunofluorescence staining was performed either by harvesting HLOs from BME-2 domes in the 24-well plate or with HLOs *in situ* on the pillar plate. In the case of BME-2 dome culture, BME-2 domes containing organoids were collected in cold dPBS^-/-^ through pipetting with wide bore tips into a 1.5 mL Eppendorf tube and left for 10 minutes to settle by gravity. The HLOs were fixed using 4% paraformaldehyde (PFA; ThermoFisher Scientific; J19943K2) for 15 minutes at room temperature by shaking five minutes intervals. PFA was diluted 4-folds with PBS to prevent organoid detachment from the pillar plate. After washing, the HLOs were permeabilized with 500 µL of 0.5% Triton X-100 (Sigma; T8787) in dPBS^-/-^ for three times at five-minute intervals. After permeabilization, HLOs were exposed to 500 µL of 5% normal donkey serum (NDS) in the permeabilization buffer for 1 hour at room temperature. For primary antibody staining, the HLOs were treated with different primary antibodies according to the manufacturer’s recommendation in 300 µL of the blocking buffer for overnight at 4°C. The HLOs were rinsed three times with PBS for 30 minutes at ten-minute intervals. The samples were incubated with appropriate secondary antibodies according to the manufacturers recommended concentration and 1 µg/mL of nuclear stain DAPI for 1 hour at room temperature. The HLOs were further washed with 1 mL of dPBS^-/-^ three times at ten-minute intervals. The HLOs were treated with 25 μL of mounting solution Visikol^®^ Histo-M™ (Visikol; HM-30) to clear the organoids. For the HLOs on the pillar plate, all the immunofluorescence staining steps were performed by sandwiching the pillar plate with a 384DeepWellPlate containing 80 µL of respective solutions and incubating the sandwiched plates under the same conditions mentioned above. Images were acquired using a fluorescence microscope (Keyence, BZ-X810). The specific names of primary and secondary antibodies used are listed in **Supplementary Table 2 and 3**, respectively.

### Measurement of the secretion of albumin and coagulation factors with ELISA assays

To measure the level of albumin and coagulation factor secretion from HLOs on the pillar plate, 80 μL of the culture medium in the 384DeepWellPlate was collected at day 20 of culture after 3 days of incubation with HLOs encapsulated in BME-2 on the pillar plate. The collected culture medium was centrifuged at 250 g for 3 minutes to remove any debris, and the resulting supernatant was assayed for albumin and coagulation factors. For albumin measurement, a human albumin ELISA kit (ThermoFisher Scientific; EHALB) was used according to the manufacturer’s instruction. For coagulation factor measurement, a factor-IX ELISA kit (Novus Biologicals; NBP2-60518) was used according to the manufacturer’s instruction.

### Measurement of bile acid transport with cholyl-lysyl-fluorescein (CLF)

To analyze the function of bile acid transport in HLOs, the HLOs were washed with dPBS^-/-^ and incubated in the advanced DMEM/F12 medium containing 5 µM cholyl-lysyl-fluorescein (AAT Bioquest Inc., CA, USA). After overnight treatment at 37°C in the CO_2_ incubator, the HLOs were rinsed with dPBS^-/-^ thrice and replaced with the advanced DMEM/F12 medium without CLF. The HLOs were then imaged by using an automated fluorescence microscope with 20x objective lens (Keyence; BZ-X800E).

### Measurement of coagulation kinetics using the HLO culture medium

Kinetic coagulation assays were performed with the day 20 HLO medium. The HLO medium without incubation with HLOs was used as a negative control. Factor V, factor IX, and factor XII deficient plasmas (Prolytix; SKU: FV-ID; SKU: FIX-ID; SKU: FXII-ID, respectively) were activated with the Siemens Healthineers Dade Actin™ FS Activate PTT Reagent (Fischer Scientific; 23-044-649) to monitor kinetic partial thromboplastin time (kPTT). Human factor VII deficient plasma (Prolytix; SKU: FVII-ID) was activated with rabbit thromboplastin (Millipore Sigma; 72162-96-0) to generate kinetic prothrombin time (kPT). For each human deficient plasma with the day 20 HLO medium collected from the 24-well plate with BME-2 domes or the pillar plate, the kPT and kPTT curves were generated and compared.

Briefly, a 96-well plate was used to perform the kPT and kPTT assays^17^. The 96-well plate was kept on ice and 10 μL of 10 mg/mL fibrinogen and 10 μL of the respective human deficient plasma were added and mixed in the individual wells. An additional 63 μL of day 20 HLO medium collected from the 24-well plate with BME-2 domes or the pillar plate was added, followed by the addition of 5 μL of Dade Actin or 1 mg/mL rabbit thromboplastin. The entire reaction was recalcified with 6 μL of 100 mmol/L CaCl_2_. The plate was then removed from the ice, wiped thoroughly, and incubated at room temperature for 4 minutes. Next, the plate was placed in a Synergy H1 plate reader (Biotek, Winooski, VT) at 37℃ and shaken for 1 minute. The absorbance at 405 nm representing fibrin formation was recorded every minute for 30 minutes. The kinetic curves were obtained by plotting the absorbance at 405 nm as a function of time.

### Statistical analysis

The statistical analysis was performed using GraphPad Prism 9.3.1 (Graph-Pad Software, San Diego, CA, USA). All the data were expressed as mean ± SD. Student’s t-test was used for comparison between two group. The statistically significant difference between the control and test groups was indicated by **** for P<0.0001, *** for p < 0.001, ** for p < 0.01, * for p < 0.05, and ns for not significant (p > 0.05).

## Results

### Generation and characterization of endodermal progenitor cells (EPCs) and mesodermal vascular progenitor cells (VPCs) from human iPSCs

Despite great advancements in organoid research recently, the lack of vasculature restricts the *in vivo*-like complexity and functionality, thereby limiting the utility of human organoids. In the precedent works, different approaches were demonstrated to generate hepatic vascular models. In 2013, Takebe *et al.* generated vascular liver buds by co-culturing cells from different sources including iPSC-derived hepatic endodermal cells, human umbilical cord endothelial cells (HUVECs), and mesenchymal stem cells^3,4^. Other studies have demonstrated liver bud formation by combining hepatic endoderm, endothelium, and septum mesenchyme generated from isogenic cell sources in a microwell plate-based platform^5,18,19^. Giuseppe *et al*. demonstrated hepatic-like vascular organoids by combining endothelial cells derived from adipose tissues with human iPSCs-derived embryoids^20^. Kim *et al.* generated multi-lineage vascular liver organoids by assembling hepatic endoderm, hepatic stellate cell-like cells, and endothelial cells derived from hPSCs in ultra-low attachment 96-well plates^21^. However, these studies did not show the passage ability of liver buds or organoids.

In the present work, miniature vHLOs have been generated by co-differentiating VPCs and EPCs on our unique pillar plate (**Figures 1 - 3**). In particular, we investigated the enhanced maturity of liver organoids by introducing vasculature, with emphasis on coagulation factors. Prior to iPSC differentiation, the quality of iPSCs was verified by measuring OCT4 and SOX2 stem cell markers (**Figure 1A**). The VPCs were generated through iPSC differentiation with BMP4 and CHIR99021 for 3 days. Characterization was performed by immunofluorescence staining for CD34 endothelial progenitor marker and CD31 endothelial marker, as well as qPCR analysis for GATA-4 mesodermal cells, along with VEGFA endothelial cell marker (**Figures 1B and 1C**). The VPCs differentiation was confirmed by the high expression of *CD34* and *GATA-4* genes and the high expression of *VEGF*A endothelial cell marker. The gene expression of *VEGFA* confirmed the characteristic endothelial signature at early formation process. In addition, endodermal progenitor cells were generated by iPSC differentiation with Activin A, Wnt3a, and R-spondin1 for 5 days, which were characterized by immunofluorescence staining of CXCR4 and FOXA2 as well as qPCR analysis of *CXCR4*, *FOXA2* and *SOX17,* and *OCT4* for stem cells (**Figures 1D and 1E**). The EPCs differentiation was confirmed by high expression of EpCAM, CXCR4, and FOXA2 proteins as well as high expression of *EPCAM*, *CXCR4*, *FOXA2*, *SOX17* genes. As a result, the expression level of *OCT4* gene was significantly decreased due to stem cell differentiation (**Figure 1E**). Akbari *et al*.^22^ demonstrated that the addition of R-Spondin1 along with Wnt3a increased the population of EpCAM^+^ endodermal progenitor cells and confirmed the formation of HLOs after EpCAM^+^ cell sorting. In the present work, we demonstrated that HLOs can be generated with the EpCAM^+^ cells directly without cell sorting. Morphological changes in vascular progenitor cells and endodermal progenitor cells are provided in **Supplementary Figure 1**.

**Figure 2.**
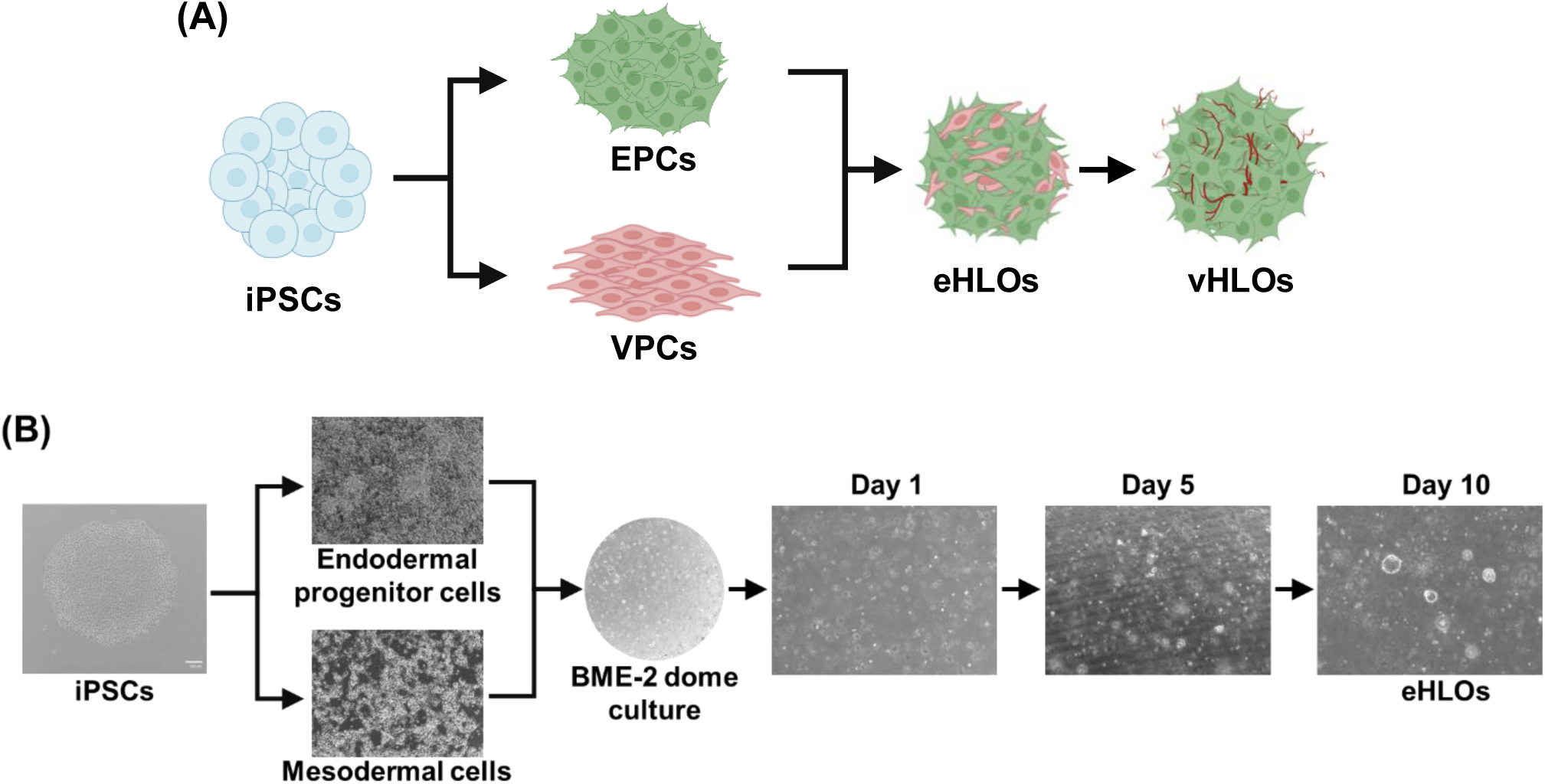
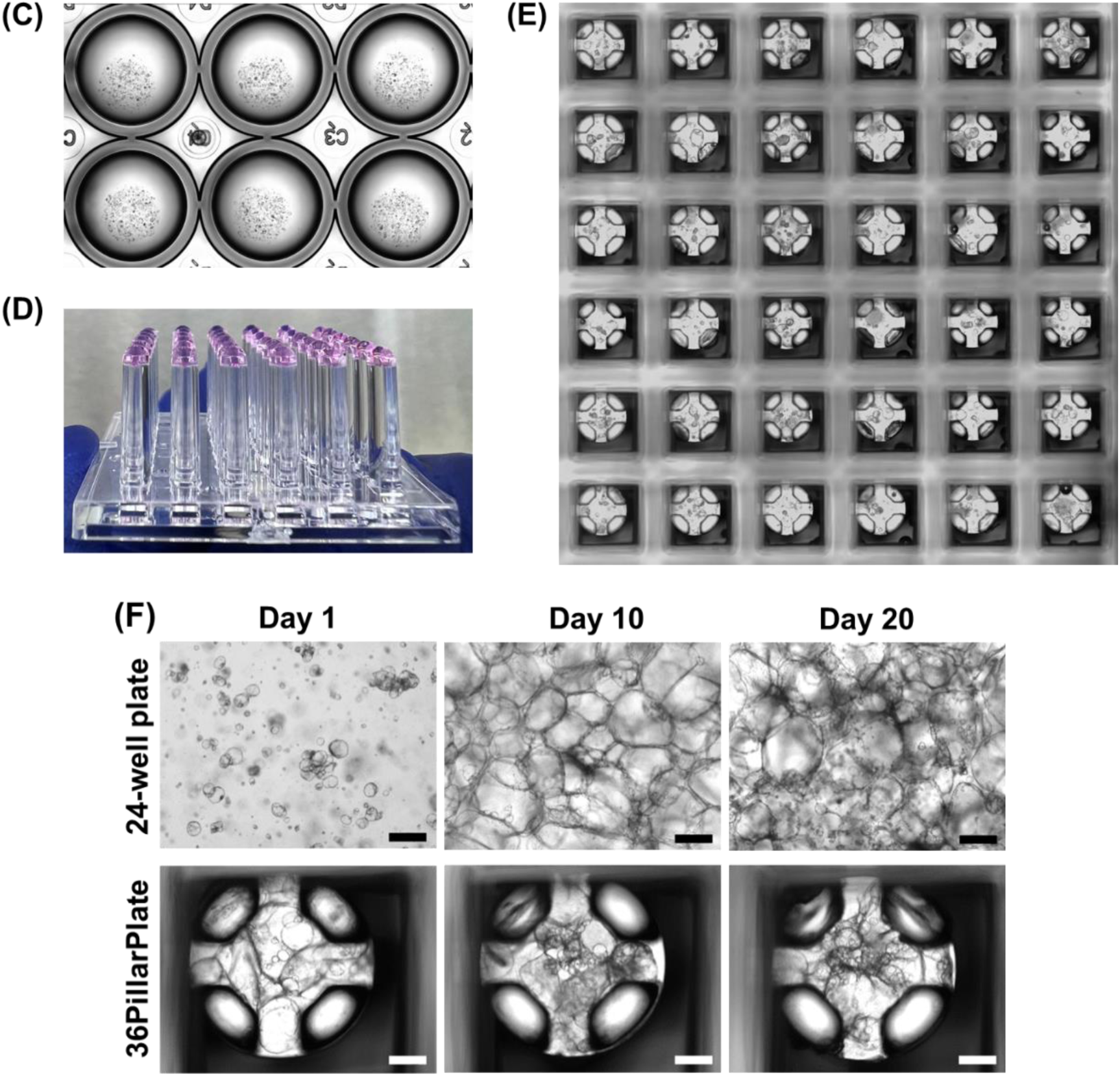
Generation of premature expandable human liver organoids (eHLOs) and mature vascular human liver organoids (vHLOs): **(A)** Schematic diagram for the generation of eHLOs by differentiation of EPCs and VPCs together in BME-2 domes in a 24-well plate and further differentiation into vHLOs. **(B)** Generation of premature eHLOs in BME-2 domes in the 24-well plate. **(C)** Picture of eHLOs in BME-2 domes in the 24-well plate. **(D)** Bioprinted eHLOs in BME-2 spots on the 36PillarPlate. **(E)** Stitched image of bioprinted eHLOs on the 36PillarPlate. **(F)** Generation of vHLOs over time in BME-2 domes in the 24-well plate and BME-2 spots on the 36PillarPlate. Scale bars: 500 μm.

**Figure 3.**
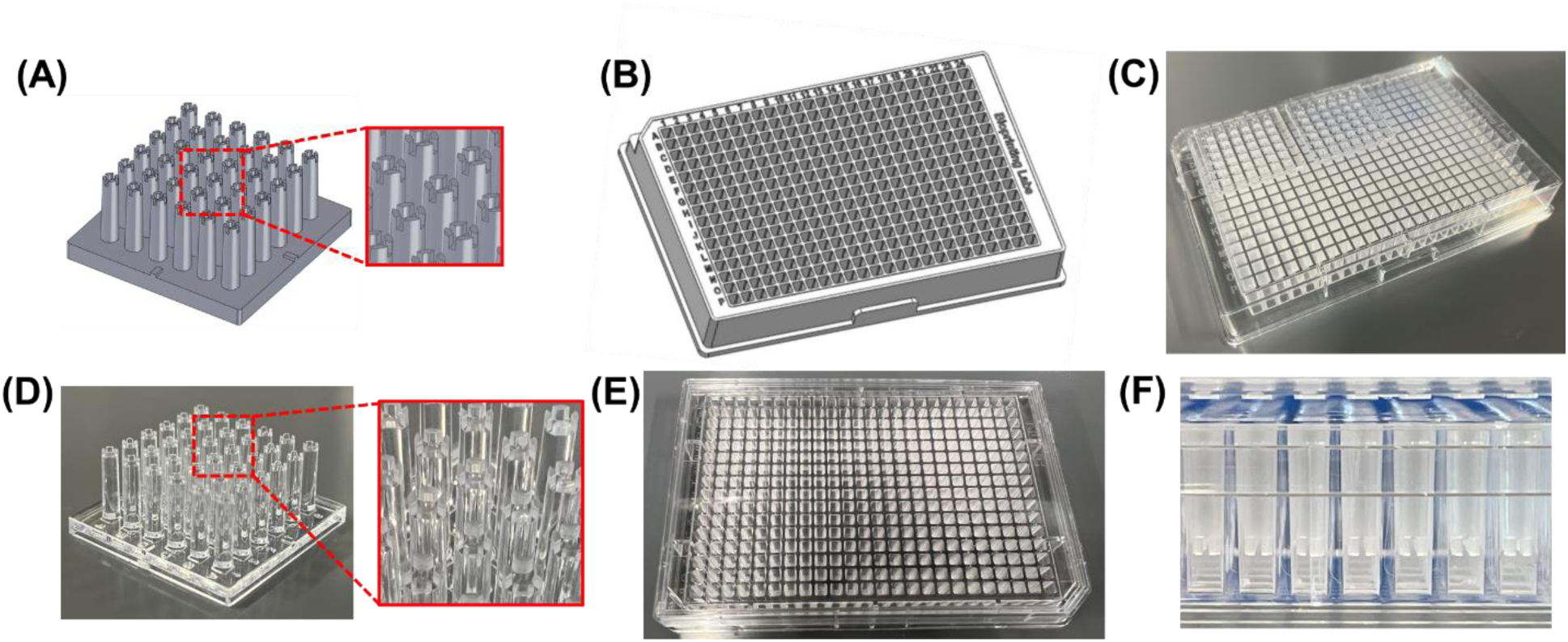
Injection-molded pillar plate and complementary deep well plate for static organoid culture. SolidWorks design of **(A)** a 36PillarPlate with a 6 x 6 array of pillars and **(B)** a 384DeepWellPlate with a 16 x 24 array of deep wells. The picture of **(C)** the 36PillarPlates sandwiched into the 384DeepWellPlate, **(D)** the 36PillarPlate and **(E)** 384DeepWellPlate manufactured by injection molding with polystyrene and **(F)** a closeup image of the pillars of the 36PillarPlate inserted into the wells of the 384DeepWellPlate for organoid culture.

### Generation of vascular human liver organoids (vHLOs)

Expandable human liver organoids (eHLOs) were generated by mixing EPCs and VPCs in BME-2 and co-differentiating them in BME-2 domes in a 24-well plate for 10 days, which were further differentiated into vHLOs for 20 days (**Figure 2**). From day 5, spheroids were formed, which gradually grew into eHLOs with 300 - 500 µm in diameter within 10 - 14 days of differentiation (**Figure 2B**). VEGF was supplemented in the eHLO formation medium to promote the endothelial specification of mesodermal cells as well as maturation of eHLOs. After 10 - 14 days of co-differentiation, eHLOs were harvested from BME-2 domes, physically dissociated into small cell clumps, mixed with fresh BME-2, and then dispensed into the 24-well plate (**Figure 2C**) or printed onto the 36PillarPlate at 5 µL per pillar (**Figures 2D and 2E and Supplementary Video 1**). We introduced a unique pillar plate with sidewalls and slits (e.g., 36PillarPlate) and a complementary deep well plate (e.g., 384DeepWellPlate) for static culture of vHLOs in the present work (**Figure 3**) as normal HLOs cultured on the pillar plate showed higher maturity as compared to their counterpart, Matrigel dome-cultured HLOs in the 24-well plate^23^. The pillar plate can be coupled with a perfusion plate to further enhance organoid maturity^24–26^.

After differentiation of vHLOs in the 24-well plate and on the pillar plate for 20 days, the morphology of vHLOs (**Figure 2F and Supplementary Videos 2 and 3**) as well as the expression level of hepatic biomarkers were extensively investigated (**Figure 4**). The morphology of vHLOs on the 36PillarPlate was closely resembled that of vHLOs cultured in BME-2 domes in the 24-well plate (**Figure 2F and Supplementary Figure 2**). Immunofluorescence staining was performed to characterize the maturity of vHLOs on the pillar plate by measuring the expression of ALB albumin, HNF4a early hepatic progenitor cells, and E-cadherin for cell-cell tight junction formation (**Figure 4A**). The maturity of vHLOs was further confirmed by measuring the high expression level of hepatic maturation genes such as *ALB* albumin as well as *CYP3A4* and *UGT1A1* drug metabolizing enzymes (**Figure 4B**), which was compared with the maturity of normal HLOs generated by using the Ouchi *et al* protocol^8^. The expression level of *EPCAM* gene was greatly reduced due to EPC differentiation into mature HLOs (**Figures 4C)**. Immunofluorescence staining also confirmed the presence of vascular networks in both eHLOs and vHLOs by measuring CD31 endothelial cell marker (**Figures 4D and 4E**). This was further confirmed by measuring the high expression level of vascular genes in vHLOs on the pillar plate, including *VEGFA*, *VWF*, and *CDH5*, as compared to those in eHLOs (**Figure 4F**). These results indicate that vascular cells were present in vHLOs and persisted through the differentiation process along with the hepatic cell population. Furthermore, immunofluorescence staining of vHLOs confirmed the high expression of collagen 1A1 (COL1A1) and collagen 3A1 (COL3A1), which was compared with eHLOs (**Figures 4G and 4H**). COL1A1 and COL3A1 are crucial hepatic extracellular matrix (ECM) proteins in the liver. Finally, the high secretion level of albumin, the HLO maturity marker, in vHLOs on the pillar plate as compared with that of eHLOs (**Figure 4I**) as well as hepatic bile acid transportation in vHLOs in BME-2 domes in the 24-well plate were demonstrated (**Figure 4J**). The fluorescent bile acid, cholyl-lysyl-fluorescein (CLF), was accumulated in all the vHLOs generated in BME-2 domes in the 24-well plate. These results signified the enhanced complexity of vHLOs generated by co-differentiating mesoderm derived vascular progenitor cells and endodermal progenitor cells in BME-2.

**Figure 4.**
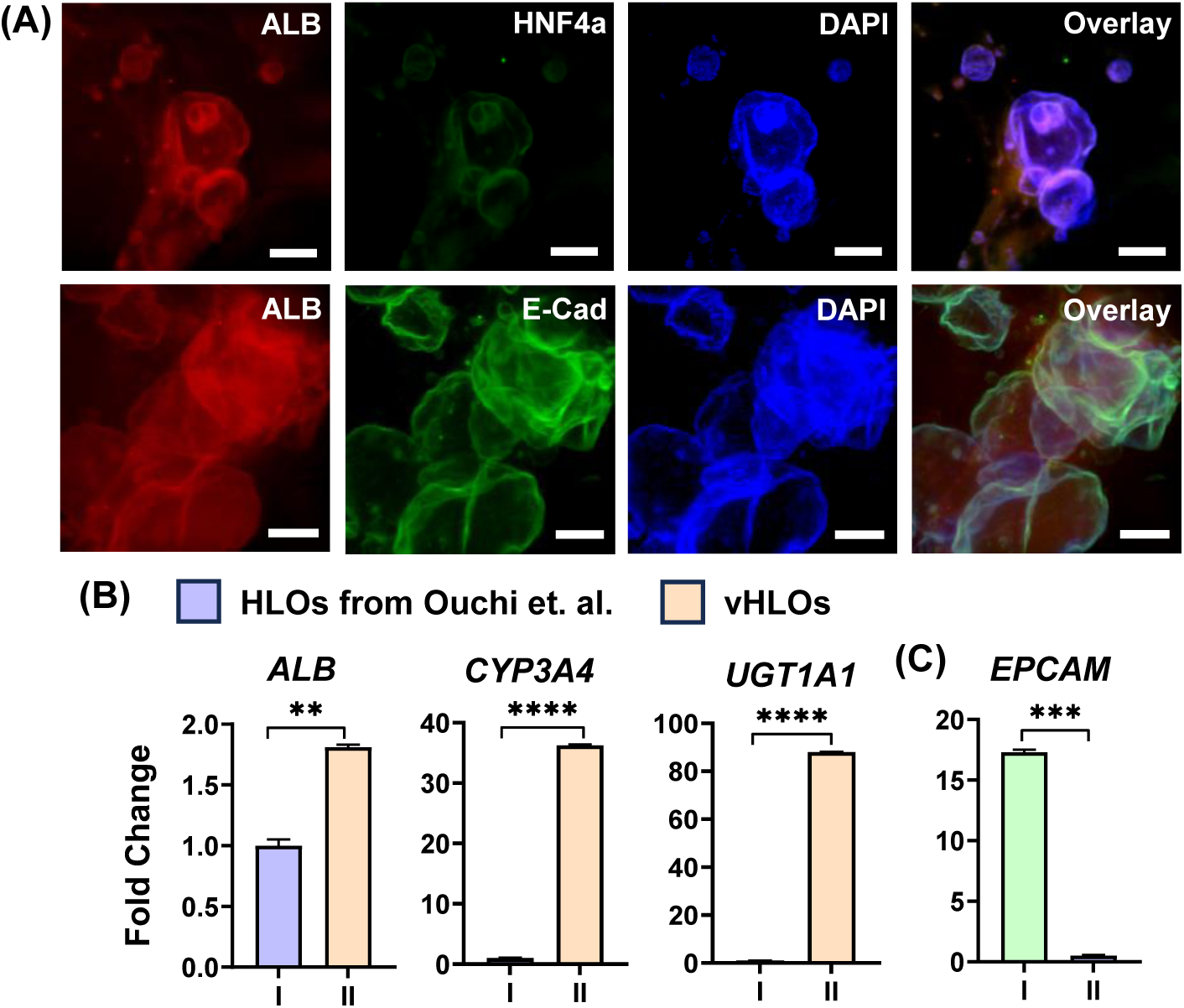
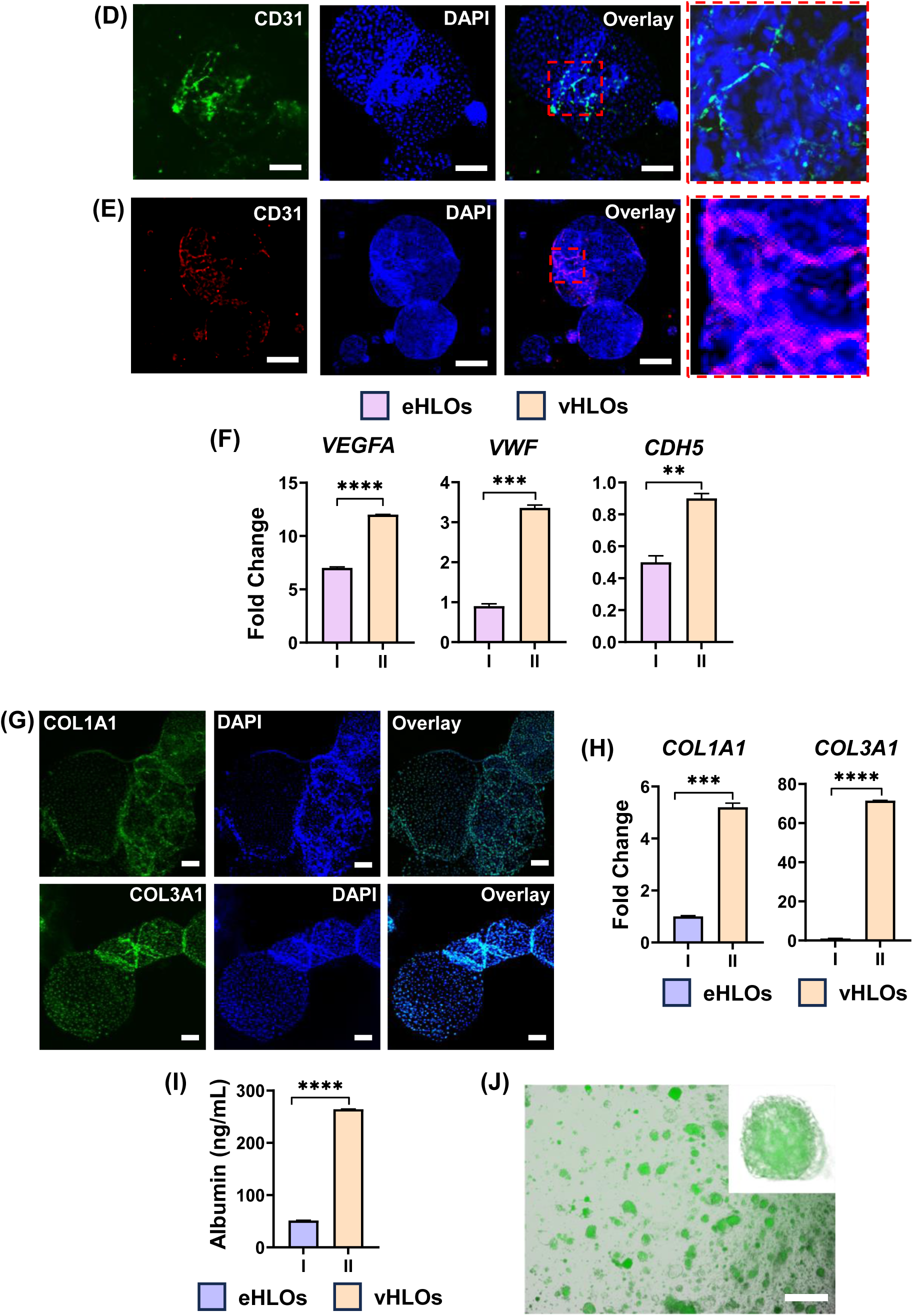
Assessment of hepatic biomarkers in vHLOs: **(A)** Immunofluorescence staining of vHLOs on the pillar plate after 20 days of differentiation for ALB albumin, HNF4a hepatic progenitor cells, and E-cadherin cell-cell tight junction formation. Scale bars: 100 µm. **(B)** The expression level of liver maturity genes including *ALB* albumin as well as *CYP3A4* and *UGT1A1* drug metabolizing enzymes, compared with *EpCAM* endodermal progenitor cells (EPCs). ‘I’ represents HLOs generated by using the Ouchi et al. protocol whereas ‘II’ represents vHLOs generated by our protocol. n = 6. **(C)** qPCR analysis of EpCAM progenitor marker from (I) EPCs and (II) matured HLOs. n = 6. The expression levels were compared with the hiPSCs. **(D)** Immunofluorescence staining of eHLOs on the the pillar plate for CD31 endothelial cell marker. Scale bars: 200 µm. **(E)** Immunofluorescence staining of vHLOs on the pillar plate for CD31 endothelial cell marker. Scale bars: 200 µm. **(F)** The expression level of vascular genes in eHLOs in the 24-well plate (n =6) and vHLOs on the pillar plate (n = 36) including *VEGFA* endothelial cells, *VWF* von Willebrand factor for making blood clotting, and *CDH5* Cadherin 5 responsible for vascular epithelium expression. The expression levels were compared with the hiPSCs. **(G)** Immunofluorescence staining of vHLOs on the pillar plate for COL1A1 and COL3A1 collagen markers. Scale bars: 100 µm. **(H)** The expression level of liver extracellular matrix (ECM) genes in eHLOs in the 24-well plate (n =6) and vHLOs on the pillar plate (n = 36) including *COL1A1* and *COL3A1* collagen. **(I)** The secretion level of albumin from eHLOs in the 24-well plate (n =6) and vHLOs on the pillar plate (n = 36). **(J)** Bile acid accumulation in vHLOs. Scale bar: 100 µm.

### Secretion of coagulation factors from vHLOs

Since the liver is the main organ generating most coagulation factors, we tested the expression level of coagulation factor genes in vHLOs (**Figure 5A**). We found that there was a significant increase in the expression levels of transcripts of *F5* (coagulation factor V), *F7* (coagulation factor VII), *F8* (coagulation factor VIII), and *F9* (coagulation factor IX) genes in vHLOs, compared to normal HLOs without vasculature. In addition, we measured the secretion level of factor IX protein from vHLOs, which was higher than that from normal HLOs (**Figure 5B**). Furthermore, the function of coagulation factors in the vHLO medium was confirmed by coagulation kinetic assays by mixing the vHLO medium from the pillar plate and the 24-well plate with coagulation factor-deficient plasma. The vHLO medium without incubation with vHLOs was used as a control. Interestingly, coagulation was induced most significantly with the vHLO medium obtained from the pillar plate for coagulation factors V, VII, IX, and XII, indicating that these coagulation factors are secreted from vHLOs. Interestingly, the vHLO medium obtained from the pillar plate exhibited higher activity for factor IX and factor XII than factor V and factor VII. In addition, the vHLO medium obtained from the 24-well plate showed higher activity of factor IX and factor XII compared to the normal HLO medium from the 24-well plate (**Supplementary Figure 4**). In all cases, the vHLO medium from the pillar plate appeared to have higher coagulation factor activities compared to that from the 24-well plate (**Figure 6B**). Furthermore, the coagulation factor activities were decreased significantly when the premature vHLO medium from day 12 vHLOs was used, although it remained slightly higher compared to the 24-well plate (**Supplementary Figure 5**).

**Figure 5.**
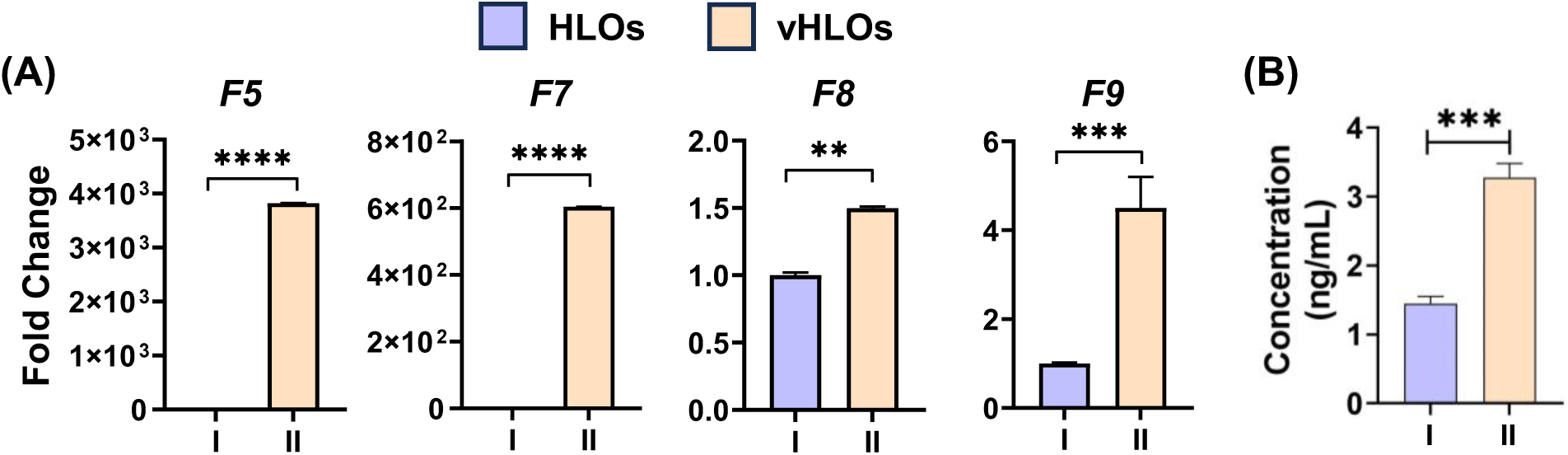
Secretion of coagulation factors from vHLOs: **(A)** The expression level of coagulation factor genes in normal HLOs in the 24-well plate (n = 6) and vHLOs on the pillar plate (n = 36) including *F5*, *F7*, *F8* and *F9*. **(B)** Secretion of factor IX from normal HLOs in the 24-well plate (n = 6) and vHLOs on the pillar plate (n = 36). The significance of gene expression difference was analyzed by Student’s t test.

**Figure 6.**
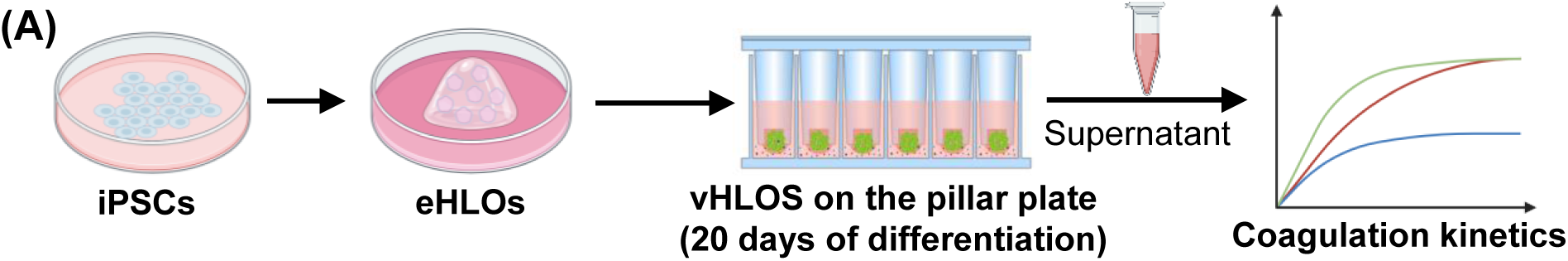
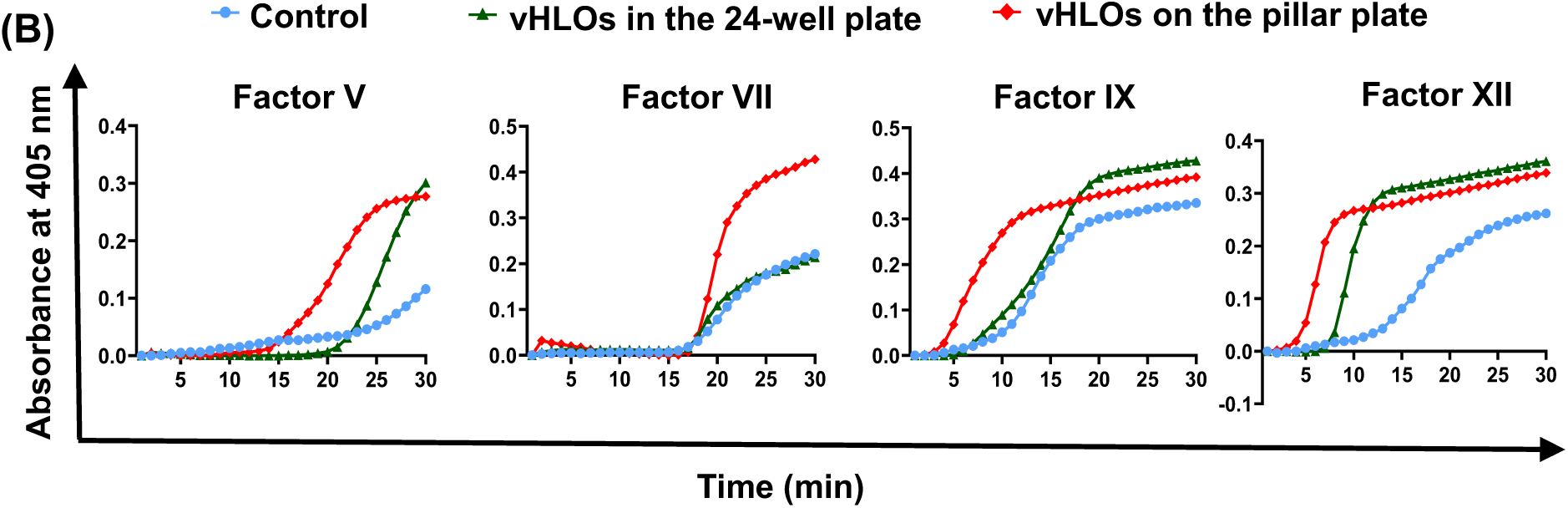
Assessment of kinetics of coagulation factors generated by vHLOs: **(A)** Schematic diagram for coagulation kinetics with coagulation factors generated by vHLOs on the pillar plate. **(B)** Comparison of coagulation kinetics with coagulation factors generated by vHLOs in the 24-well plate and on the pillar plate. The advanced DMEM/F12 medium without incubation with vHLOs was used as a control.

## Discussion

In this study, vHLOs were generated by coordinated differentiating endodermal progenitor cells and mesodermal vascular progenitor cells obtained from single iPSC line together. It has been reported that vascular cells are originated from mesoderm^16,27^, while liver morphogenesis occurs from endoderm^28,29^. We generated EpCAM^+^ endodermal progenitor cells by iPSC differentiation with Activin A, Wnt3a, and R-spondin as reported by Akbari et al.^22^. Mesodermal vascular progenitor cells were generated by iPSC differentiation with BMP4 and CHIR99021, which were further differentiated into endothelial cells by the addition of VEGF, along with HLO formation. Our approach resulted in significantly higher level of coagulation factor secretion than the published protocols by using the combination of the early-stage progenitor cells, highlighting the essential role of vascular mesoderm in HLO maturation and coagulation factor secretion. The expression level of *VEGFA*, a key marker of endothelial cells, was high in both eHLOs and vHLOs (**Figure 4F**), indicating vasculature maturation within the liver organoids. The addition of VEGFA directed differentiation of vascular progenitor cells into mature CD31^+^ endothelial cells (**Figure 4D**). The expression of vascular transcriptional and translational markers confirmed the sustained development of vasculature within the vHLOs throughout the expansion and maturation stages.

During the vHLO maturation process, the organoids became denser in structure, and BME-2 gradually disappeared, suggesting the initiation of their own ECM secretion. Under microscopic observation, vHLOs resembled liver lobules like structures (**Supplementary Figure 2**). In addition, the maturity of vHLOs generated on the pillar plate was higher with high coagulation factor secretion compared to that of vHLOs in BME-2 domes in the 24-well plate (**Figure 6B**). This result suggests that miniaturization of vHLO culture on the pillar plate provides better availability of growth factors and nutrients during long-term organoid differentiation and growth compared to conventional BME-2 dome culture. In addition, this implies that the choice of cell culture systems could play a crucial role in organoid maturation. With our pillar plate, we were able to generate organoids with small volume of cell culture media (80 µL/well or pillar), making it technically more feasible for downstream applications. In addition, we previously demonstrated that cell death in the core of spheroids can be reduced, and the maturity of brain and liver organoids can be enhanced greatly by simply differentiating cells in the pillar/perfusion plate and facilitating rapid diffusion of nutrients and oxygen^24–26,30^.

Thrombosis, the formation of blood clots within blood vessels, is a major contributor to global disease mortality and morbidity^31^. This condition primarily arises from alterations in blood composition, blood flow, and vessel wall integrity^32^. The liver plays a crucial role in blood coagulation by generating various coagulation factors and anticoagulants that regulate platelet function, fibrinolysis, and thrombopoietin production^33,34^. The endothelium, the innermost single layer of endothelial cells lining the blood vessels, is imperative in regulating blood fluidity and providing an antithrombotic surface. The endothelium produces several factors such as nitric oxide, prostacyclin, Von Willebrand factor (vWF), thrombomodulin, and endothelin, which modulate platelet reactivity, coagulation, fibrinolysis, and vascular contractility^35^. Therefore, endothelial cells are pivotal in modulating thrombosis, making them a potential target for regulating thrombosis^35,36^. The coagulation cascade involves two distinct pathways: the intrinsic and extrinsic pathways^37^. The subendothelial matrix activates the extrinsic pathway initiated by tissue factors (TFs), located on subendothelial cells. In contrast, the intrinsic pathway includes factors XII, XI, IX, and VIII. Both pathways converge on the activation of factor X (the common pathway) through a series of interactions with other factors, forming the prothrombinase complex, which converts prothrombin to thrombin. Thrombin further strengthens the platelet plug, preventing additional bleeding, by cleaving fibrinogen to generate fibrin, which polymerizes to form a stable fibrin clot^38,39^. To mimic such conditions *in vitro* and develop 3D tissue models, it is essential to mimic organogenesis by incorporating the mesoderm-derived endothelial cells and the endoderm-derived hepatic cell population in the coagulation process.

The coagulation cascade involves a series of interactions among various coagulation factors from both intrinsic and extrinsic pathways^37^. *In vivo*, the coagulation system is activated by factor III (tissue factor) present on the surface of subendothelial cells and culminates with the activation of factor X for clot formation^40,41^. Our vHLOs expressed and secreted a critical coagulation factor VII for the extrinsic pathway, and the intrinsic pathway factors VIII, IX, and XII, and the common pathway cofactor V (**Figures 5 and 6**). All these factors contribute to thrombotic events *in vivo*. In addition, vWF and collagens were also expressed in vHLOs (**Figure 4**). Collagens are exposed and attached at the site of vessel injury in the circulatory system, while vWF mediates the platelet adherence, serving as primary stimuli *in vivo*^40,42^. These findings indicate that vHLOs exhibited relevant thrombotic signatures seen *in vivo*. The cellular crosstalk between endothelial cells and liver cells, as well as coagulation factor secretion by vHLOs, could closely mimic the physiological events in the native liver.

The liver is the primary organ to produce most of the coagulation factors except vWF, factor III, and factor VIII^43,44^. The endothelial cells-derived vWF is a large multimeric protein that plays an essential role in hemostasis and thrombosis^45^. Previous reports have shown the secretion of coagulation factors from iPSC-derived, hepatocyte-like cells from different sources and species^46–49^. Later, the focus shifted to 3D cell models. Giuseppe *et al*.^20^ demonstrated the secretion of coagulation factors by combining endothelial cells derived from adipose tissues with human iPSC-derived embryoids and differentiating them into hepatic-like vascular organoids. Takebe *et al.* reported the secretion of coagulation factor VIII from liver buds generated by combining hepatic endodermal, endothelial, and mesenchymal cells^1^. Another report showcased the secretion of coagulation factors from vascularized liver organoids generated in a bioreactor, utilizing 125 mL media to culture the organoids^50^.

Currently, zebrafish, rodent, and swine models are used for thrombosis modeling^40^. However, these animal models may provide different physiology and genetic makeups from humans^12,51^. Our vHLOs cultured on the pillar plate could be coupled with microphysiological systems may further improve their physiological relevance and clinically translatable outcomes. In addition, the pillar plate could be used for organoid-based assays *in situ*, including scale-up production of organoids, compound testing, as well as organoid staining and imaging.

### Conclusions

The vHLOs, generated by merging endodermal and mesodermal progenitor cells, demonstrated advanced maturation when bioprinted onto a pillar plate compared to a 24-well plate dome culture. These vHLOs showed a more pronounced expression of essential hepatic biomarkers than HLOs derived from published protocols. Moreover, the vHLOs secreted hepatocyte-specific coagulation factors V, VII, IX, and XII, as well as endothelial cell-derived factors like vWF and factor VIII. Thus, the vHLO models could be invaluable for investigating liver formation from combined endoderm and mesoderm, offering a contrast to HLOs that originate from a single endoderm layer. Additionally, the integration of our user-friendly, high-throughput pillar plate platform with vHLOs promises to improve disease modeling and drug testing efficiency while reducing costs.

## Data availability

The authors declare that all the data supporting the results of this study are available within the article and its Supplementary Information.

## Acknowledgments

This study was financially supported by the National Institutes of Health (NIEHS R43ES035653, NCATS R44TR003491, NIDDK UH3DK119982, and NHLBI R01HL159399).

## Author contributions

V.K.R.L.: Conceived, designed, and conducted the organoid experiments, interpreted, and analyzed the results, and wrote the manuscript.

S.S.: Conducted the control organoid experiments with the previously published protocol and wrote the manuscript with related information.

A.A.Q.: Conducted the thrombosis experiments, analyzed results, and contributed to manuscript writing.

S.D.: Conducted the thrombosis experiments and contributed to manuscript writing.

P.A.: Helped the organoid experiments and data analysis.

A.R.: Helped manuscript writing and proofreading.

P.J.: Designed the thrombosis experiments, interpreted the results, revised the manuscript, and supervised the thrombosis research.

M.Y.L.: Conceived and designed the pillar/perfusion plates, planned the organoid experiments, interpreted the results, wrote, and revised the manuscript, and supervised the entire project.

## Supplementary information

**Supplementary Table 1.**
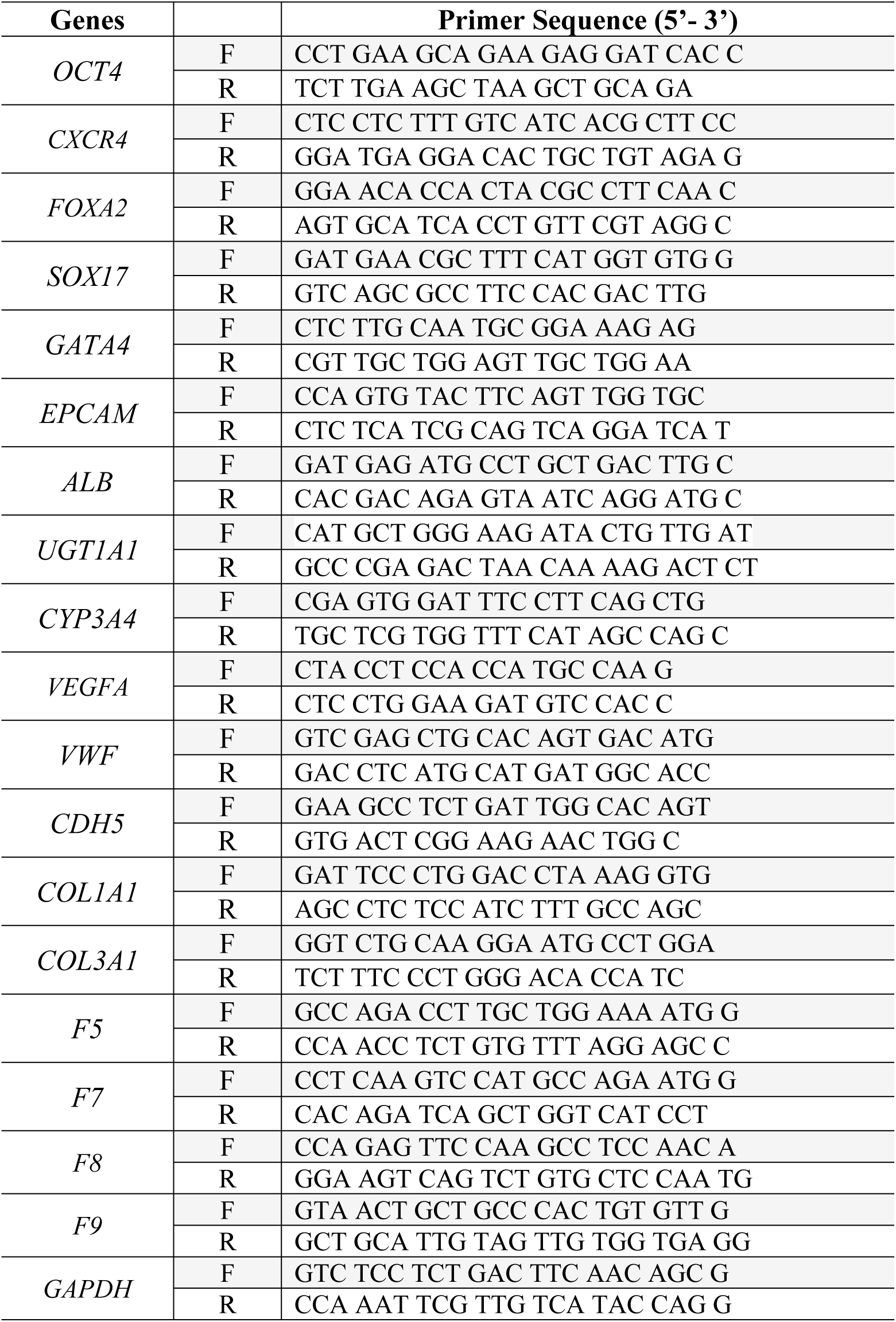
List of primers.

**Supplementary Table 2.** List of primary antibodies.

| **Primary antibodies** | **Host/Isotype** | **Vendor** | **Dilution/Working concentration** |
| --- | --- | --- | --- |
| OCT3/4 | Mouse/monoclonal (IgG<sub>2</sub>) | sc-5279<br>SantaCruz | 1:40 |
| SOX2 | Mouse/monoclonal (IgG<sub>1</sub>) | sc-365823<br>SantaCruz | 1:40 |
| CXCR4 | Mouse/monoclonal (IgG<sub>1</sub>) | sc-53534<br>SantaCruz | 1:40 |
| HNF3β/FOXA2 | Goat/polyclonal (IgG) | AF2400<br>R&D Systems | 5 μg/mL |
| HNF4a | Mouse/monoclonal (IgG<sub>1</sub>) | sc-374229<br>SantaCruz | 1:40 |
| ALB | Rabbit/polyclonal (IgG) | ab2406<br>Abcam | 1:100 |
| E-cad | Mouse/monoclonal (IgG<sub>1</sub>) | sc-21791<br>SantaCruz | 1:40 |
| EpCAM | Mouse/monoclonal (IgG<sub>1</sub>) | sc-25308<br>SantaCruz | 1:40 |
| CD31 | Mouse/monoclonal (IgG<sub>1</sub>) | 3528S<br>Cell Signaling<br>Technology | 5 μg/mL |
| CD34 | Mouse/monoclonal | 11034942<br>Invitrogen | 5 μg/mL |
| COL1A1 | Rabbit/polyclonal (IgG) | NBP130054<br>Novus | 5 μg/mL |
| COL3A1 | Rabbit/polyclonal (IgG) | NB600594<br>Novus | 5 μg/mL |

**Supplementary Table 3.** List of secondary antibodies.

| Secondary antibodies | Host | Vendor | Dilution |
| --- | --- | --- | --- |
| Donkey anti-Mouse IgG (H+L) Highly Cross-Adsorbed Secondary Antibody, Alexa Fluor™ 488 | Donkey | A-21202<br>Invitrogen | 1:200 |
| Donkey anti-Mouse IgG (H+L) Highly Cross-Adsorbed Secondary Antibody, Alexa Fluor™ 647 | Donkey | A-31571<br>Invitrogen | 1:200 |
| Donkey anti-Rabbit IgG (H+L) Highly Cross-Adsorbed Secondary Antibody, Alexa Fluor 488 | Donkey | A-21206<br>Invitrogen | 1:200 |
| Donkey anti-Rabbit IgG (H+L) Highly Cross-Adsorbed Secondary Antibody, Alexa Fluor 647 | Donkey | A-31573<br>Invitrogen | 1:200 |
| Donkey anti-Goat IgG (H+L) Highly Cross-Adsorbed Secondary Antibody, Alexa Fluor 647 | Donkey | A-21447<br>Invitrogen | 1:200 |

**Supplementary Figure 1.**
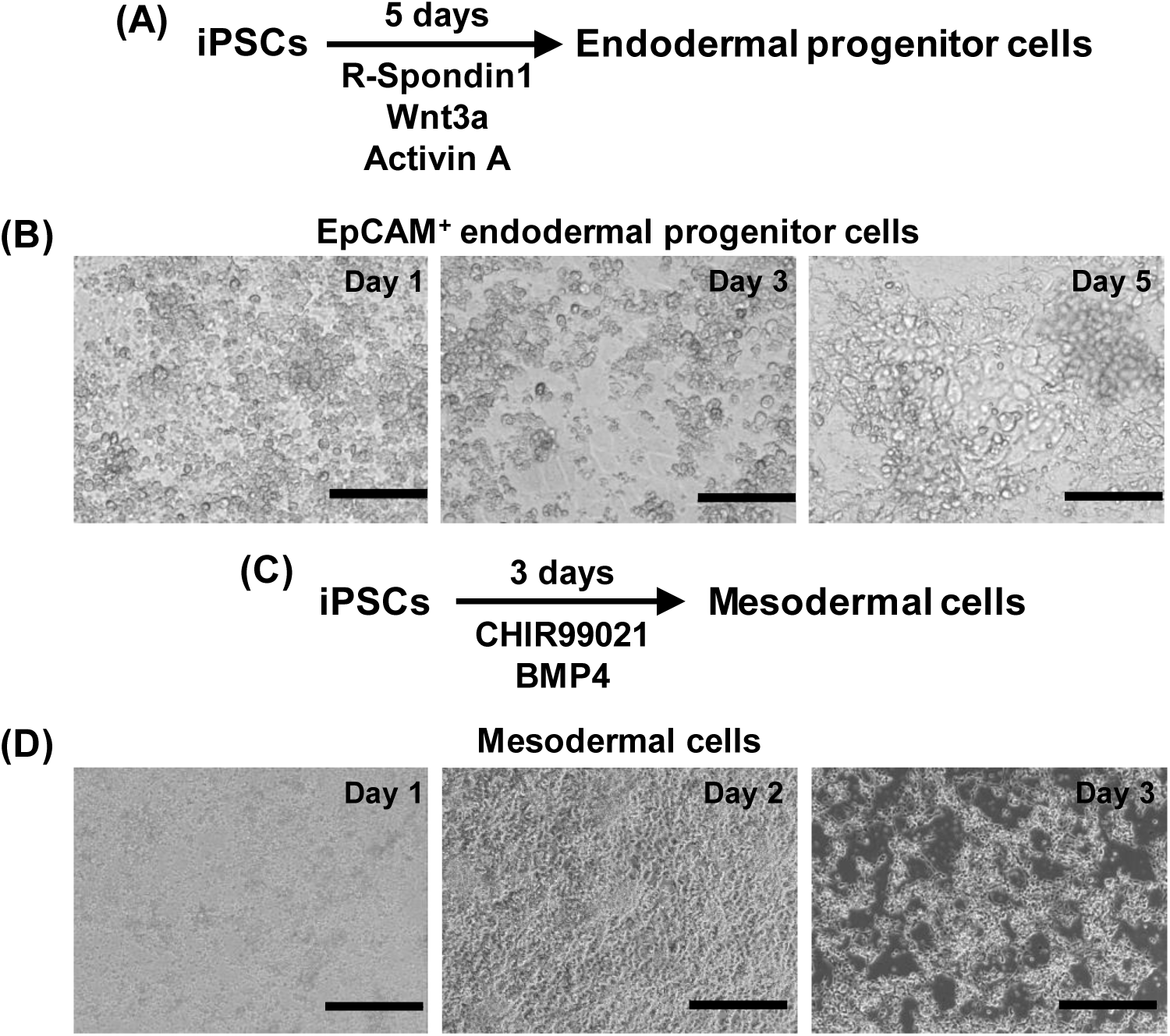
**(A)** Schematic diagram for the differentiation of endodermal progenitor cells from human iPSCs for 5 days in the 6-well plate. **(B)** Changes in cell morphology over time for endodermal progenitor cells. **(C)** Schematic diagram for the differentiation of mesodermal cells from human iPSCs for 3 days in the 6-well plate. **(D)** Changes in cell morphology over time for mesodermal cells. Scale bars: 100 µm.

**Supplementary Figure 2.**
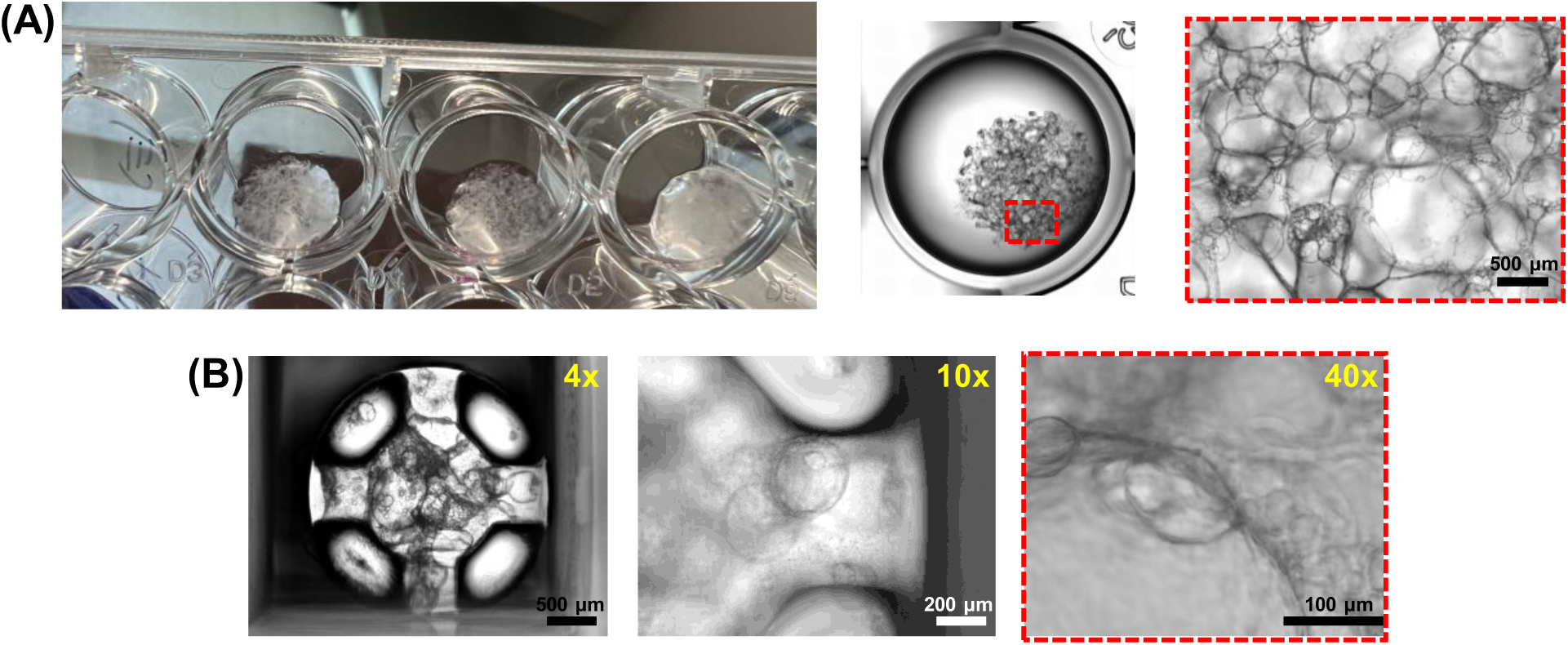
**(A)** The differentiation of vHLOs in the 24-well plate for 20 days. High-density vHLOs were observed with naked eyes in 24-wells. BME-2 started to degrade over time, which was replaced with ECMs generated by vHLOs. **(B)** The differentiation of vHLOs on the pillar plate for 20 days. Scale bars: 500, 200, and 100 µm. Similar morphology was observed on the pillar plate.

**Supplementary Figure 3.**
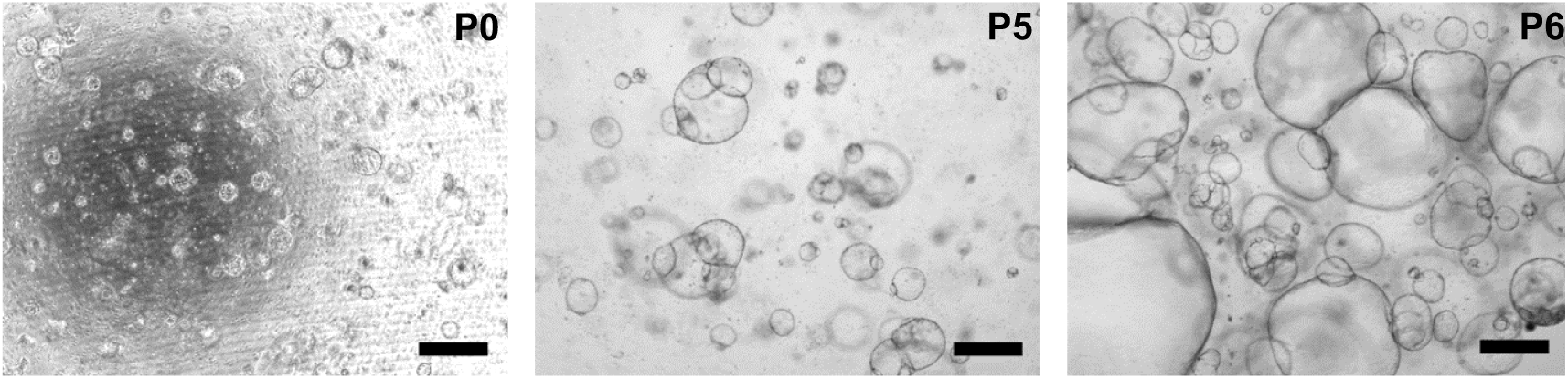
Passage of eHLOs in BME-2. After 5 - 7 days of culture, eHLOs were physically dissociated with a pipette tip and seeded in fresh BME-2 for passage and maturation studies. Scale bars: 500 µm.

**Supplementary Figure 4.**
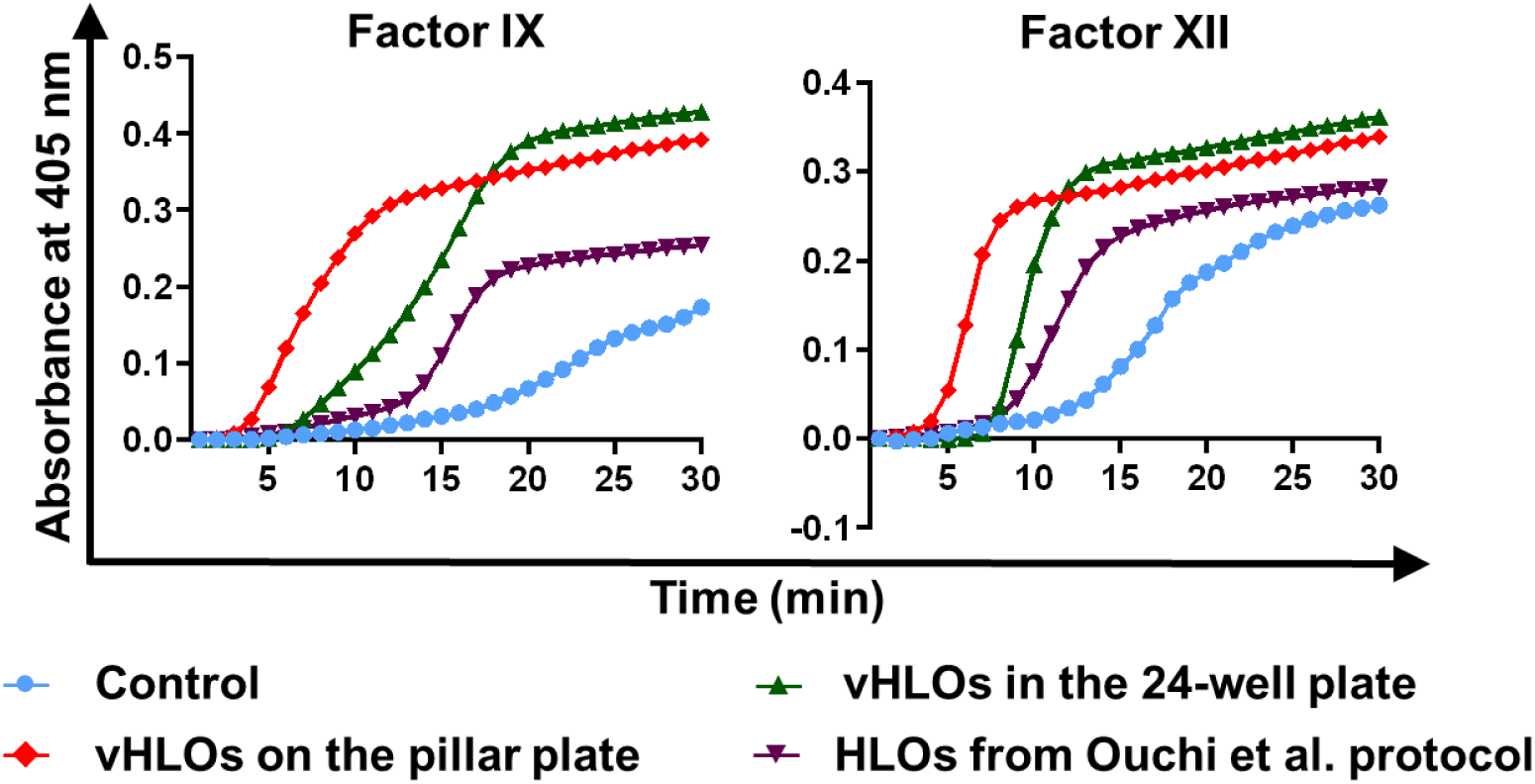
Comparison of coagulation kinetics with coagulation factors generated by day 20 vHLOs in the 24-well plate and on the pillar plate. Normal HLOs were generated by using the Ouchi et al protocol for comparison. The advanced DMEM/F12 medium without incubation with vHLOs was used as a control. Both factor IX and factor XII showed higher sensitivity with vHLOs from the pillar plate, indicating higher maturity.

**Supplementary Figure 5.**
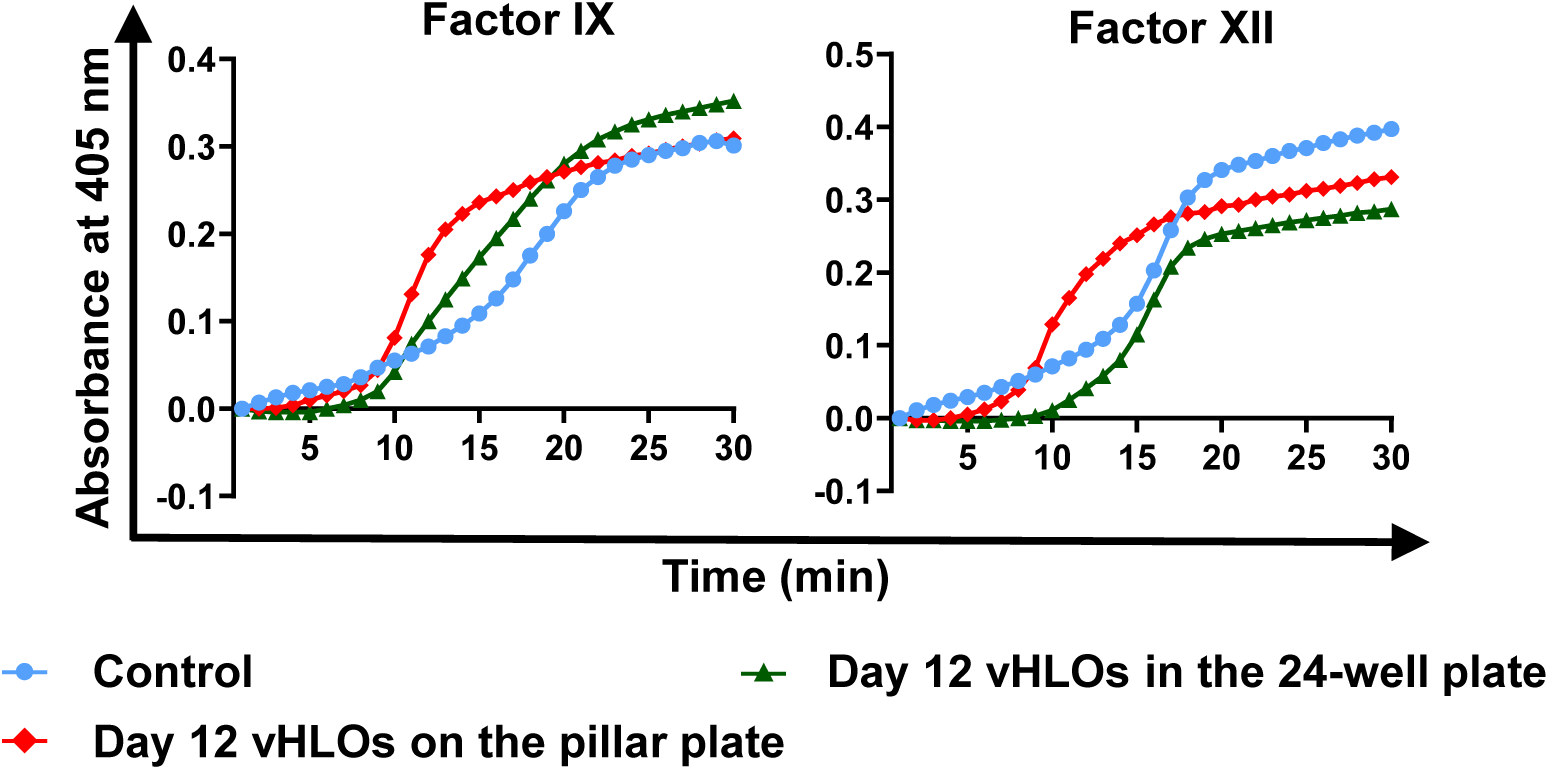
Comparison of coagulation kinetics with coagulation factors generated by day 12 vHLOs in the 24-well plate and on the pillar plate. The advanced DMEM/F12 medium without incubation with vHLOs was used as a control. The amount of factor IX and factor XII generated by day 12 vHLOs was insufficient to induce fast coagulation, indicating immaturity of day 12 vHLOs.

## Supplementary video clips

**Video 1.** Bioprinting of eHLOs on the 36PillarPlate.

https://youtu.be/ZIwA3I8iU80

**Video 2.** vHLOs on the 36PillarPlate after 20 days of differentiation. The video was created with the Z-stacking images.

https://youtu.be/94YFiSfk5MM

**Video 3.** vHLOs in the 24-well plate after 20 days of differentiation. The video was created with the Z-stacking images.

https://youtu.be/-F7C_CVJCm8

